# Chondrocyte-specific RUNX2 Overexpression Causes Chondrodysplasia During Development, but is Not Sufficient to Induce OA-like Articular Cartilage Degeneration in Adult Mice Without Injury

**DOI:** 10.1101/470005

**Authors:** Sarah E. Catheline, Donna Hoak, Martin Chang, John P. Ketz, Matthew J. Hilton, Michael J. Zuscik, Jennifer H. Jonason

## Abstract

RUNX2 is a transcription factor critical for chondrocyte maturation and normal endochondral bone formation. It promotes the expression of factors catabolic to the cartilage extracellular matrix and is shown to be upregulated in human osteoarthritic cartilage and in murine articular cartilage following joint injury. To date, *in vivo* studies of RUNX2 overexpression in cartilage have been limited to forced expression in osteochondroprogenitor cells preventing investigation into the effects of chondrocyte-specific RUNX2 overexpression during development or in postnatal articular cartilage. Here, we used the *Rosa26^Runx2^* allele in combination with the inducible *Col2a1^CreERT2^* transgene or the inducible *Acan^CreERT2^* knock-in allele to achieve chondrocyte-specific RUNX2 overexpression (OE) during embryonic development or in the postnatal articular cartilage of adult mice, respectively. RUNX2 OE was induced at E13.5 for all developmental studies and resulted in a phenotype resembling chondrodysplasia at E18.5. Histology and *in situ* hybridization analyses suggest an early onset of chondrocyte hypertrophy and accelerated terminal maturation in the limbs of the RUNX2 OE embryos compared to control embryos. Additionally, RUNX2 OE resulted in enhanced TUNEL staining indicative of increased chondrocyte apoptosis throughout all regions of the growth plate. For all postnatal studies, RUNX2 OE was induced at 2 months of age. Surprisingly, no histopathological signs of OA or cartilage catabolism were observed even six months following induction of RUNX2 OE in postnatal animals. Using the meniscal/ligamentous injury (MLI), a surgical model of knee joint destabilization and meniscal injury, however, we found that chondrocyte-specific RUNX2 OE accelerates the progression of OA pathogenesis following joint trauma. Histomorphometry and OARSI scoring confirmed decreased cartilage area two months following injury in the RUNX2 OE joints compared to control joints. Further, the numbers of MMP13-positive and TUNEL-positive chondrocytes were significantly greater in the articular cartilage of the RUNX2 OE joints compared to control joints one month following injury. Collectively, our data support that RUNX2 OE in growth plate chondrocytes is sufficient to promote their hypertrophy and terminal maturation during development. While RUNX2 overexpression alone is surprisingly insufficient to induce catabolic changes to the postnatal articular cartilage, it can accelerate the progression of post-traumatic OA. These results suggest that although increased RUNX2 expression may predetermine the rate of OA onset and/or progression following traumatic joint injury, alone this change is not sufficient to initiate the OA degenerative process.

## INTRODUCTION

Endochondral ossification is a carefully orchestrated process regulated by a number of growth factors, signaling pathways, and transcription factors. It is responsible for development of the long bones and driven by a process of progressive cell differentiation. Initially, mesenchymal progenitor cells condense and differentiate into committed chondrocytes, whose gene expression is largely controlled by the master chondrogenic transcription factor SOX9^(1-3)^. Committed chondrocytes then begin to rapidly proliferate, building rudimentary skeletal elements rich in ACAN and COL2A1. The cells at the very center of these elements eventually exit the cell cycle and become pre-hypertrophic, expressing *Runx2* and *Ihh*^(4-7)^. These cells further mature into hypertrophic chondrocytes marked by their expression of *Col10a1* as well as other factors that promote extracellular matrix (ECM) mineralization and vascular invasion. Finally, late stage hypertrophic chondrocytes express *Mmp13.* These cells eventually undergo apoptosis, or differentiate into cells of the osteoblast lineage, leaving behind a cartilaginous template upon which bone is formed and the primary ossification center established^(8-11)^.

Postnatally, secondary ossification centers form in the epiphyses to separate the transient growth plate cartilage from the permanent articular cartilage layer that lines the ends of the long bones. In contrast to the highly proliferative growth plate chondrocytes, articular chondrocytes at homeostasis are largely non-proliferative^(12,13)^. Their matrix is composed of distinct zones that vary both biochemically in composition and cellularly with respect to chondrocyte size and gene expression, resembling the layers seen in growth plate cartilage during endochondral bone formation. The superficial zone cells along the surface of the articular cartilage have a flattened morphology and express *Prg4,* which encodes LUBRICIN to promote joint lubrication^(14,15)^. The middle zone (transitional/radial zone) cells have a rounded morphology and express matrix components COL2A1 and ACAN. The deep zone cells most closely resemble hypertrophic chondrocytes and express COL10A1 and MMP13. The middle and deep zones are separated by a basophilic line called the tidemark; this line segments the unmineralized and mineralized portions of the articular cartilage^(16)^. Despite the fact that the zones of the articular cartilage are reminiscent of those in the growth plate cartilage, articular chondrocytes of the superficial or middle zones do not enter hypertrophy or undergo terminal maturation at homeostasis. However, it has been reported that during the development of osteoarthritis (OA), these articular chondrocytes begin to mimic growth plate chondrocytes and aberrantly express proteins associated with chondrocyte hypertrophy, such as COL10A1 and MMP13^(17-20)^. Whether onset of hypertrophy alone is sufficient to promote OA pathogenesis, though, is unclear.

RUNX2 (also known as CBFA1) is a transcription factor that plays a critical role in induction of chondrocyte hypertrophy and terminal maturation as well as in osteoblast differentiation during endochondral bone formation. RUNX2 is expressed by both prehypertrophic and hypertrophic chondrocytes, as well as by cells within the perichondrium^(4,6,21,22)^. In chondrocytes, direct target genes of RUNX2 include *Ihh*, *Col10a1,* and *Mmp13*, all markers of chondrocyte hypertrophy and maturation^(23-26)^. Global deletion of *Runx2* in mice causes a loss or severe delay of chondrocyte hypertrophy and a complete lack of bone formation throughout the skeletons of RUNX2-deficient embryos^(21,22,27)^. Interestingly, *Runx2* deletion in osteochondroprogenitor cells leads to a long bone and vertebral phenotype that is similar to that seen in global knockouts, whereas deletion of *Runx2* from committed osteoblasts results in no skeletal abnormalities^(28,29)^. Further, studies of RUNX2 overexpression using *Col2a1*-driven transgenes lead to accelerated and ectopic endochondral ossification due to enhanced chondrocyte hypertrophy and terminal differentiation^(6)^. Overall, these results emphasize the importance of RUNX2 in chondrocytes for normal endochondral bone development.

Based on the well-documented role of RUNX2 in chondrocyte maturation, several studies have examined the postnatal role of RUNX2 in articular cartilage, and in the induction of OA. While baseline RUNX2 expression in the articular cartilage is minimal, it is induced following traumatic joint injury, along with its downstream targets COL10A1 and MMP13^(19,30,31)^. Global RUNX2 haploinsufficiency as well as chondrocyte-specific *Runx2* deletion were both shown to be chondroprotective following traumatic knee joint injury^(30,32)^. Expression of matrix degrading enzymes and overall cartilage degeneration were decreased in both models suggesting that RUNX2 expression in chondrocytes contributes to the pace of OA progression. Whether RUNX2 overexpression is sufficient to induce postnatal OA onset, however, has not been able to be addressed by previous genetic models due to perinatal lethality.

In this study, we aimed to explore the ability of chondrocyte-specific RUNX2 overexpression to affect chondrocyte maturation during development and within the postnatal articular cartilage. We used the *Rosa26^Runx2^* allele, previously shown to rescue bone formation in *Runx2^-/-^* embryos carrying the *Col2a1^Cre^* transgene^(33)^, in combination with the inducible *Col2a1^CreERT2^* transgene^(34)^ or the inducible *Acan^CreERT2^* knock-in allele^(35)^ to force expression of RUNX2 specifically in immature chondrocytes during development or in postnatal cartilage, respectively. Using these genetic models, we found that chondrocyte-specific RUNX2 overexpression resulted in chondrodysplasia of developing embryos, while RUNX2 overexpression in the articular chondrocytes of adult mice surprisingly did not induce changes to the articular cartilage at homeostasis. Following traumatic joint injury, however, RUNX2 overexpression resulted in more severe articular cartilage degeneration likely due to increased expression of MMP13 and enhanced apoptosis of articular chondrocytes. These data suggest that while increased expression of RUNX2 alone is not sufficient to promote cartilage degeneration, it can accelerate the development of post-traumatic OA following joint injury.

## MATERIALS AND METHODS

### Mice

Animal studies used protocols approved by the University of Rochester Committee on Animal Resources. Mice were housed in a room using microisolator technology and kept at 70°F-73°F. They had *ad libitum* access to food and water at all times. *Col2a1^CreERT2^*, *Acan^CreERT2^*, and *Rosa26^Runx2^* (*R26^Runx2^*) mice were previously described ^(33-35)^. Female *R26^Runx2/Runx2^* mice were bred with male *Col2a1^CreERT2/+^; R26^Runx2/+^* or *Acan^CreERT2^; R26^Runx2/+^* mice to generate experimental mice heterozygous or homozygous for the *R26^Runx2^* allele in combination with either the *Col2a1^CreERT2^* transgene *Acan^CreERT2^* knock-in allele. Cre-negative littermate mice were used as controls. To induce RUNX2 overexpression (OE) in embryos, pregnant females were administered tamoxifen once at E13.5 (0.1 mg/g body weight, IP). For postnatal and MLI studies, 2-month-old mice were administered tamoxifen for 5 consecutive days (0.1 mg/g body weight, IP). For the injury studies, meniscal/ligamentous injury (MLI) was performed as previously described on Cre-negative control (Control) and *Acan^CreERT2/+^; R26^Runx2/+^* (RUNX2 OE) mice at 10 weeks of age ^(36)^. Briefly, a 5-mm incision was made on the medial side of the joint; the medial collateral ligament (MCL) was then transected and the medial meniscus detached at the anterior tibial attachment site. Contralateral sham joints only received the initial incision. Numbers of mice per experimental group are as indicated: E14.5 *Col2a1^CreERT2^* for histology (Cre^-^, n = 7; Cre^+^ *R26^Runx2+/-^*, n = 6; Cre^+^ *R26^Runx2/Runx2^*, n = 4), E18.5 *Col2a1^CreERT2^* for skeletal preparations (Cre^-^, n = 4, Cre^+^ *R26^Runx2+/-^*, n = 7; Cre^+^ *R26^Runx2/Runx2^*, n = 6), E18.5 *Col2a1^CreERT2^* for histology (Cre^-^, n = 6; Cre^+^ *R26^Runx2+/-^*, n = 3; Cre^+^ *R26^Runx2/Runx2^*, n = 9), 3-month-old *Acan^CreERT2^* (Cre^-^, n = 5, Cre^+^; *R26^Runx2+/-^*, n = 4), 8-month-old *Acan^CreERT2^* (Cre^-^, n = 9; Cre^+^ *R26^Runx2/+^*, n = 6; Cre^+^ *R26^Runx2/Runx2^*, n = 5), 3.5-month-old *Acan^CreERT2^* with MLI at 2.5 months (Cre^-^ males, n = 6; Cre^+^ males, n = 6; Cre^-^ females, n = 6; Cre^+^ females, n = 6), 4.5-month-old *Acan^CreERT2^* with MLI at 2.5 months (Cre^-^ male, n = 6; Cre^+^ male, n = 7; Cre^-^ female, n = 6; Cre^+^ female, n = 6).

### Skeletal preparations, histology, *in situ* hybridization, and immunohistochemistry

Whole mount Alcian blue/Alizarin red skeletal staining of E18.5 embryos was performed as described^(37)^. For histology, embryonic tissues were fixed in 10% neutral buffered formalin (NBF) for 24 hours followed by decalcification in 14% EDTA (pH 7.3-7.4) for 24 hours. Adult hindlimbs were cleaned of soft tissue and fixed in 10% NBF for 3 days followed by decalcification for 1 week in 14% EDTA (pH 7.4-7.6). All fixation and decalcification steps were carried out at room temperature. Tissues were then processed for paraffin embedding and sectioning at 5 microns with postnatal hindlimbs oriented sagittally. Embryonic tissue sections were stained with Alcian blue Hematoxylin/Orange G and adult tissue sections with Safranin O/Fast green to visualize cartilage, bone, and soft tissues. *In situ* hybridization of embryonic tissue sections was performed as previously described using ^35^S-labeled riboprobes to *Col10a1* and *Mmp13*^(38,39)^. After developing, slides were counterstained with Toluidine blue. Final images were prepared by merging brightfield images with red pseudo-colored darkfield images.

For MMP13, COL10A1, and RUNX2 immunohistochemistry, sections were deparaffinized and rehydrated. Antigen retrieval was performed in 4 mg/ml pepsin solution made in 0.01 N HCl for 10 minutes at 37°C (for MMP13 and COL10A1) and in Antigen Unmasking Solution, Citrate Based (Vector Labs) overnight at 65°C (for RUNX2). Following rinses with deionized water, sections were incubated with BLOXALL to quench endogenous peroxidases (Vector Labs) for 10 minutes and rinsed in deionized water. Sections were then blocked in 2.5% normal horse serum for 30 minutes and incubated with primary antibody for 1 hour at room temp (anti-COL10A1 diluted 1:500, Clone X53, Quartett) or overnight at 4°C (anti-MMP13 diluted to 4 μg/ml, ab75606, Abcam; anti-RUNX2 diluted to 0.125 μg/ml, HPA022040, Sigma). Sections were rinsed in phosphate buffered saline + 0.1% tween-20 (PBST) and incubated with ImmPress HRP anti-rabbit IgG peroxidase polymer secondary (anti-MMP13, Vector Labs) or ImmPress HRP anti-mouse IgG peroxidase polymer secondary (anti-COL10A1, Vector Labs) for 30 minutes. After subsequent rinses in PBST and deionized water, COL10A1 or MMP13 were detected by DAB peroxidase HRP substrate (Vector Labs). For RUNX2, the ImmPress Excel HRP anti-rabbit IgG peroxidase polymer secondary kit was used per manufacturer’s instructions (Vector Labs). All slides were counterstained using Mayer’s hematoxylin. For negative control slides for anti-MMP13 and anti-RUNX2 staining, rabbit IgG (Vector Labs) was used at a concentration of 4 μg/ml (MMP13) or 0.125 μg/ml (RUNX2). As the stock concentration of COL10A1 is unknown, negative control slides were incubated with secondary antibody only. TUNEL staining was performed per manufacturer’s instructions using the *In Situ* Cell Death Detection Kit, Fluorescein (Sigma Roche).

### Human cartilage samples

As previously described^(40)^, a tissue microarray was created using 2 to 3 cores from 11 normal cartilage samples and 14 injured cartilage samples. An IRB-approved protocol was used to collect discarded cartilage tissue from orthopaedic surgery patients. Normal cartilage was collected from hip fracture patients. Knee articular cartilage was collected from patients undergoing arthroscopic surgery 4 weeks following meniscal injury. No identifiers are associated with the tissues. Tissue was fixed, decalcified, and processed for embedding into paraffin.

### Chondrocyte isolation and culture

Primary chondrocytes from the sterna and ribs of P3 mice were isolated as described^(41)^ and cultured in DMEM (Life Technologies) containing 10% FBS and 1% penicillin/streptomycin (Life Technologies). Ad5-CMV-GFP and Ad5-CMV-Cre adenoviruses were purchased from Baylor Vector Development Laboratory (Houston, TX) and used to infect the chondrocytes at a MOI of 1000. Forty-eight hours following infection, adenovirus was replaced with standard culture media supplemented with 50 μg/ml ascorbic acid. After an additional 48 hours, cells were either fixed with 4% paraformaldehyde and incubated with 1-Step NBT/BCIP Solution (Thermo Fisher Scientific) for detection of Alkaline Phosphatase or harvested for Western blotting or quantitative reverse transcription PCR.

### Western blotting

Western blotting was performed as described previously^(40)^. Primary antibodies were used as follows: mouse anti-RUNX2 (1:500, D130-3, MBL) and mouse anti-β-ACTIN (1:5000, AC-74, Sigma-Aldrich).

### Isolation of total RNA from mouse articular cartilage

Using scalpels, articular cartilage was shaved from the tibial plateau and frozen over dry ice. Using the Bullet Blender Gold (Next Advance), the cartilage was homogenized in 250 μl of TRIzol (Thermo Fisher) with a mix of 2, 1, and 0.5 mm RNase-free zirconium oxide beads (Next Advance) for 10 minutes. Phases were separated using 100 μl chloroform per TRIzol manufacturer’s instructions (Thermo Fisher), and the aqueous layer and one volume of 70% ethanol were transferred to a RNeasy MinElute Spin Column from the RNeasy Micro Kit (QIAGEN). mRNA was then isolated per Micro Kit instructions (QIAGEN).

### Quantitative reverse transcription PCR

Total RNA was isolated from cell cultures using the RNeasy Mini Kit (Qiagen) per manufacturer’s instructions or from mouse tibial cartilage as described above. RNA was reverse-transcribed into cDNA using the iScript cDNA Synthesis Kit (Bio-Rad). Real-time PCR was performed using a Rotor-Gene Q real-time PCR cycler (Qiagen) and the PerfeCTa SYBR Green SuperMix (Quanta Biosciences) according to manufacturer’s instructions. Primer sequences used are as follows (5’-3’): *β-actin* forward: AGATGTGGATCAGCAAGCAG; *β-actin* reverse: GCGCAAGTTAGGTTTTGTCA; *Runx2* forward: TGATGACACTGCCACCTCTGACTT; *Runx2* reverse: ATGAAATGCTTGGGAACTGCCTGG. *Col10a1* forward: CTTTGTGTGCCTTTCAATCG; *Col10a1* reverse: GTGAGGTACAGCCTACCAGTTTT; *Mmp13* forward: GATGACCTGTCTGAGGAAG *Mmp13* reverse: ATCAGACCAGACCTTGAAG; *Alpl* forward: TGACCTTCTCTCCTCCATCC; *Alpl* reverse: CTTCCTGGGAGTCTCATCCT

### Modified OARSI Scoring

Semi-quantitative scoring was performed on MLI knee joint sections from mice 1 and 2-months following MLI surgery using the grading system described previously^(42)^. Briefly, 3 different sections from distinct levels per MLI sample were stained with Safranin O/Fast green and randomized for scoring. Four independent scorers performed modified OARSI scoring and scores were averaged for each slide. The average score for the 3 slides from each sample were then averaged to obtain a final score for that sample.

### Histomorphometry

Histomorphometry was performed using Visiopharm image analysis software (Version 6.7.0.2590). The measure tool was used to measure limb lengths and an average limb length was calculated from 3-5 embryos per genotype. An image analysis application was designed to measure the area of mineralized (below the tidemark) and unmineralized (above the tidemark) articular cartilage. The measure tool was used as a ruler to measure the same length across the articular surface on each section, and the ROI tool was used to manually circle the relevant cartilage areas within the length measured. The application was then trained to recognize the difference between Safranin O-positive regions (denoted as SafO+) and negative, Fast green, regions (denoted as background) of the cartilage areas using the threshold method. This was done using the Contrast Red-Green feature; areas were marked SafO+ if they met a threshold of −10 pixel intensity or above, and were marked as background if they were below the −10 value. We then applied several post-processing corrections to allow the software to recognize the white interior of chondrocytes not stained by Safranin O as portions of the SafO+ area, rather than background; the change by area feature was used to change any area labeled background 50 μm^2^ or smaller that was completely surrounded by SafO+ area to SafO+. A second change by area feature was used to change any area labeled SafO+ 50 μm^2^ or smaller that was completely surrounded by background to background; this allowed Visiopharm to recognize hematoxylin-stained chondrocyte nuclei as a part of the SafO+ area. The software was then able to calculate the area of the manually circled unmineralized and mineralized cartilage, as well as the SafO+ area within those regions. Values were averaged from three sections for both sham and MLI samples, and the MLI sample normalized to the sham for each animal.

For quantification of the COL10A1+ area, an image analysis application was designed to determine the amount of tissue stained brown with DAB (denoted as DAB+) within the articular cartilage. As in the SafO application, the measure tool was used, followed by the ROI tool to circle the articular cartilage area within the length measured. The application was then trained to recognize the difference between DAB+ brown tissue and hematoxylin stained tissue (denoted as background) using the threshold method. This was done using the HDAB-DAB feature; areas were marked DAB+ if they were at or below this threshold. The software was then able to calculate the circled cartilage area, as well as the COL10A1+ area within this region.

For MMP13+ cell counting, the stereology function of Visiopharm was used to manually determine the number of MMP13+ cells and total cells. For TUNEL+ cell counting, an image analysis application was designed on Visiopharm to recognize and count TUNEL+ stained cells and FITC+ stained cells within the articular cartilage using the threshold method. As described above, the measure tool was used followed by the ROI tool to manually circle the articular cartilage area within the length measured. The method was programmed to recognize DAPI+ and FITC+ cells using the DAPI W and FITC features. Areas were marked DAPI+ if they had DAPI values at 3000 or above and FITC values lower than 900; areas were marked FITC+ if they had FITC values at or over 900. All areas with DAPI values below 3000 or FITC values below 900 were marked as background. We then applied several post-processing corrections to allow the software to exclude any background positive smaller than the smallest typical area of a cell found within the articular cartilage (less than 10 μm^2^); using the change by area feature, both DAPI+ and FITC+ areas smaller than 10 μm^2^ were changed to background. The software was then able to calculate the number of DAPI+ and FITC+ areas within the circled ROI; each positive area was counted as one cell.

### Statistics

Data are presented as the mean ± SEM. Statistical significance was determined by Student’s *t* tests or one-way ANOVA followed by Dunnett’s multiple comparisons test as indicated; *p* values less than 0.05 were considered significant.

## RESULTS

### Chondrocyte-specific RUNX2 overexpression during development leads to chondrodysplasia

To evaluate the effects of chondrocyte-specific RUNX2 overexpression on skeletal development, we generated *Col2a1^CreERT2/+^* embryos^(34)^ with one or two copies of the *R26^Runx2^* allele^(33)^ (*Col2a1^CreERT2/+^; R26^Runx2/+^* and *Col2a1^CreERT2/+^; R26^Runx2/Runx2^*, respectively). This allele allows expression of the MASNS isoform of RUNX2 in a Cre-inducible manner. Recombination was induced via administration of tamoxifen at E13.5 and embryos were harvested at E18.5. The RUNX2-overexpressing (OE) embryos had shortened trunks, limbs, and domed skulls with protruding tongues, consistent with chondrodysplasia. Whole mount Alcian blue and Alizarin red staining revealed shorter axial skeletons, smaller rib cages, and shorter forelimbs and hindlimbs (**Fig. 1A**). Analysis of histological sections revealed decreases in the total length of the femora, tibiae, and humeri of the RUNX2 OE embryos compared to their Cre-negative control littermates (**Fig. 1B**). Additionally, RUNX2 OE tibiae were bowed with increased endocortical bone (**Supplemental Fig. 1A**). Histology also showed that the columnar and hypertrophic zones of the RUNX2 OE embryos are hypocellular and disorganized compared to control littermates with some chondrocytes in the hypertrophic zone appearing abnormally large (**Fig. 1C**). Immunohistochemistry (IHC) for RUNX2 on histological sections confirmed RUNX2 overexpression in the mutant limbs compared to control limbs (**Supplemental Fig. 1B**).

**Fig. 1.**
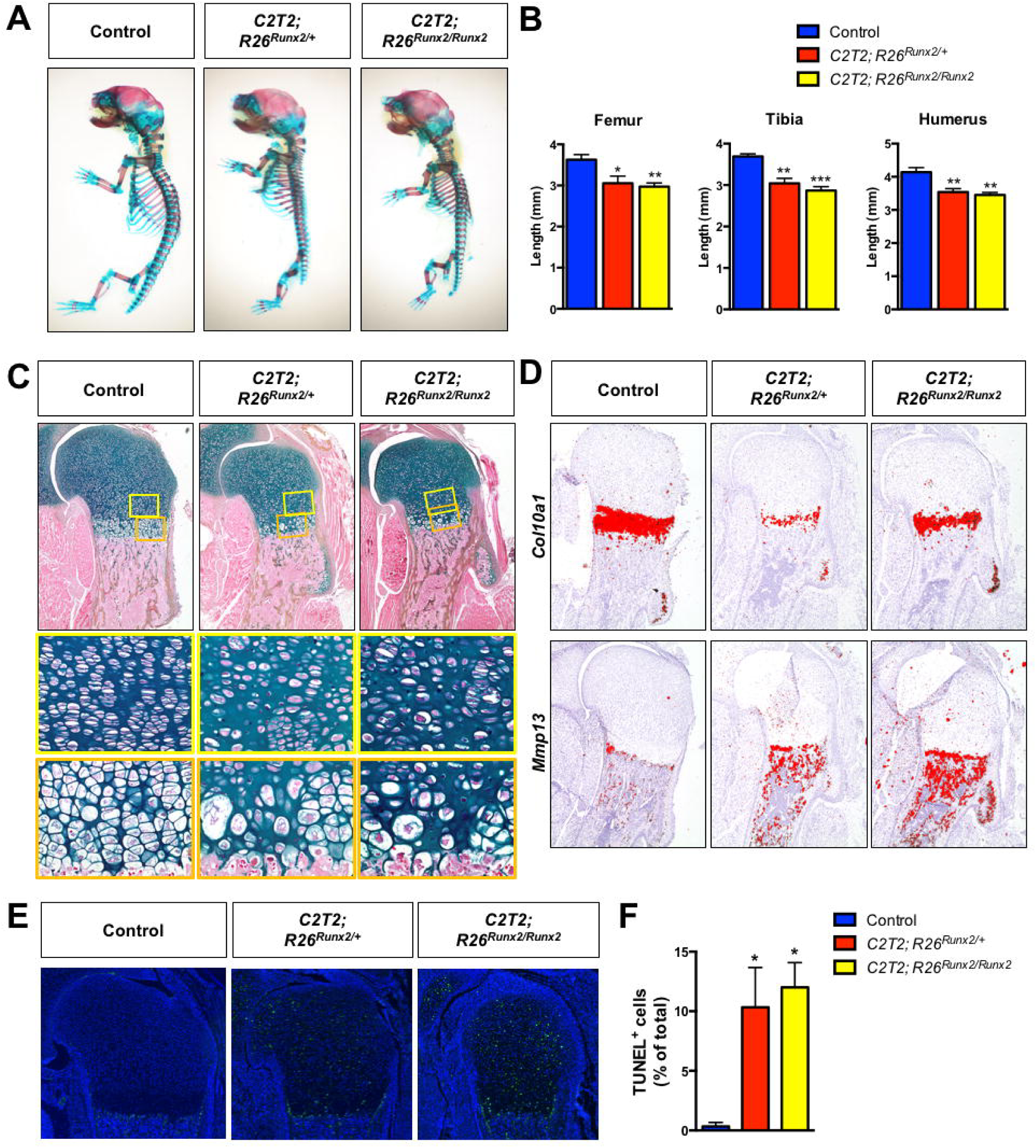
Chondrocyte-specific RUNX2 overexpression during development leads to chondrodysplasia. (A) Alcian blue/Alizarin red skeletal staining of E18.5 *R26^Runx2/+^* (Control), *Col2a1^CreERT2/+^; R26^Runx2/+^* (*C2T2; R26^Runx2/+^*), and *Col2a1^CreERT2/+^; R26^Runx2/Runx2^* (*C2T2; R26^Runx2/Runx2^*) embryos. (B) Femur (left), tibia (middle), and humerus (right) lengths from E18.5 Control, *C2T2; R26^Runx2/+^*, and *C2T2; R26^Runx2/Runx2^* embryos (n = 3 to 5 for all groups). \**p* < 0.05, \*\**p* < 0.01, \*\*\**p* < 0.001, one-way ANOVA followed by Dunnett’s multiple comparisons test. (C) Alcian blue Hematoxylin/Orange G staining of humerus growth plate sections from E18.5 Control, *C2T2; R26^Runx2/+^*, and *C2T2; R26^Runx2/Runx2^* embryos. Top panels are 5X images of the proximal humerus; middle panels are high magnification images (20X) of the proliferating and pre-hypertrophic zones (yellow boxes); bottom panels are high magnification images (20X) of the hypertrophic zone (orange boxes). (D) *In situ* hybridization for *Col10a1* (top panels) and *Mmp13* (bottom panels) on E18.5 Control, *C2T2; R26^Runx2/+^*, and *C2T2; R26^Runx2/Runx2^* proximal humerus sections. (E) TUNEL staining of proximal humerus sections from E18.5 Control, *C2T2; R26^Runx2/+^*, and *C2T2; R26^Runx2/Runx2^* embryos (5X images). (B) Quantification of TUNEL^+^ cell number as percentage of total cell number within the growth plate of the proximal humeri of E18.5 Control, *C2T2; R26^Runx2/+^*, and *C2T2; R26^Runx2/Runx2^* embryos (n = 3 for all groups). \**p* < 0.05, one-way ANOVA followed by Dunnett’s multiple comparisons test.

*In situ* hybridization analyses on E18.5 humerus sections showed that the expression domain of *Col10a1* relative to total limb length is unchanged in the RUNX2 OE embryos compared to littermate control embryos, while that of *Mmp13* is slightly expanded and maintained in the trabecular bone of the RUNX2 OE embryos (**Fig. 1D**). Analyses of RUNX2 OE embryos at E14.5, however, revealed an expanded *Col10a1* expression domain and early appearance of *Mmp13* expression relative to control embryos (**Supplemental Fig. 1C**), suggesting early onset of hypertrophy and accelerated maturation to late stages of chondrocyte hypertrophy. Primary sternal chondrocytes from *R26^Runx2/Runx2^* mice were infected with adenovirus encoding GFP or Cre recombinase and RUNX2 overexpression was confirmed at both the mRNA and protein levels (**Supplemental Fig. 2A, C**). Increased Alkaline phosphatase staining and *Col10a1*, *Alpl*, and *Mmp13* gene expression in these cultures provide *in vitro* support that the *R26^Runx2-^*expressing chondrocytes undergo accelerated hypertrophy and terminal maturation in a cell autonomous manner.

Finally, to investigate whether the hypocellularity in the E18.5 RUNX2 OE limbs could be due to cell death, we performed TUNEL staining on tissue sections. TUNEL staining was visibly increased throughout all regions of the growth plate in the RUNX2 OE limbs and quantification of TUNEL^+^ cells relative to total cell numbers supported this observation (**Fig. 1E, F**).

### Postnatal chondrocyte-specific RUNX2 overexpression alone is not sufficient to induce articular cartilage degeneration

RUNX2 expression is reported to be upregulated in human OA cartilage and in the articular cartilage of mice following traumatic joint injury^(30,32,43)^. Using a human tissue microarray, we also find evidence of increased RUNX2 expression in the articular cartilage of patients just one month following meniscal tear (**Fig. 2**). Since it is well established that joint injury increases the risk for development of OA^(44,45)^, we decided to examine whether RUNX2 overexpression alone could be sufficient to induce catabolic changes to the joint consistent with the onset of OA. Specifically, we generated *Acan^CreERT2/+^* mice with one or two copies of the *R26^Runx2^* allele (*Acan^CreERT2/+^; R26^Runx2/+^* and *Acan^CreERT2/+^; R26^Runx2/Runx2^*, respectively). Recombination was induced via administration of tamoxifen at 2 months of age, a time point following the majority of rapid skeletal growth that occurs during early postnatal life. One week following tamoxifen induction, there were no overt changes in cartilage phenotype, but RUNX2 IHC did show increased RUNX2 expression within the articular cartilage of RUNX2 OE mice (**Fig. 3A, B**). Additionally, quantitative RT-PCR from mRNA isolated from the tibial articular cartilage confirmed this finding, showing a significant increase in *Runx2* expression in the RUNX2 OE group (**Fig. 3C**). Surprisingly, at 8 months of age (6 months following tamoxifen induction), histology reveals that there are still no overt phenotypic changes to the articular cartilage of the RUNX2 OE mice relative to control mice, despite evidence that RUNX2 is still expressed in the articular chondrocytes at this time (**Fig. 3D, E**).

**Fig. 2.**
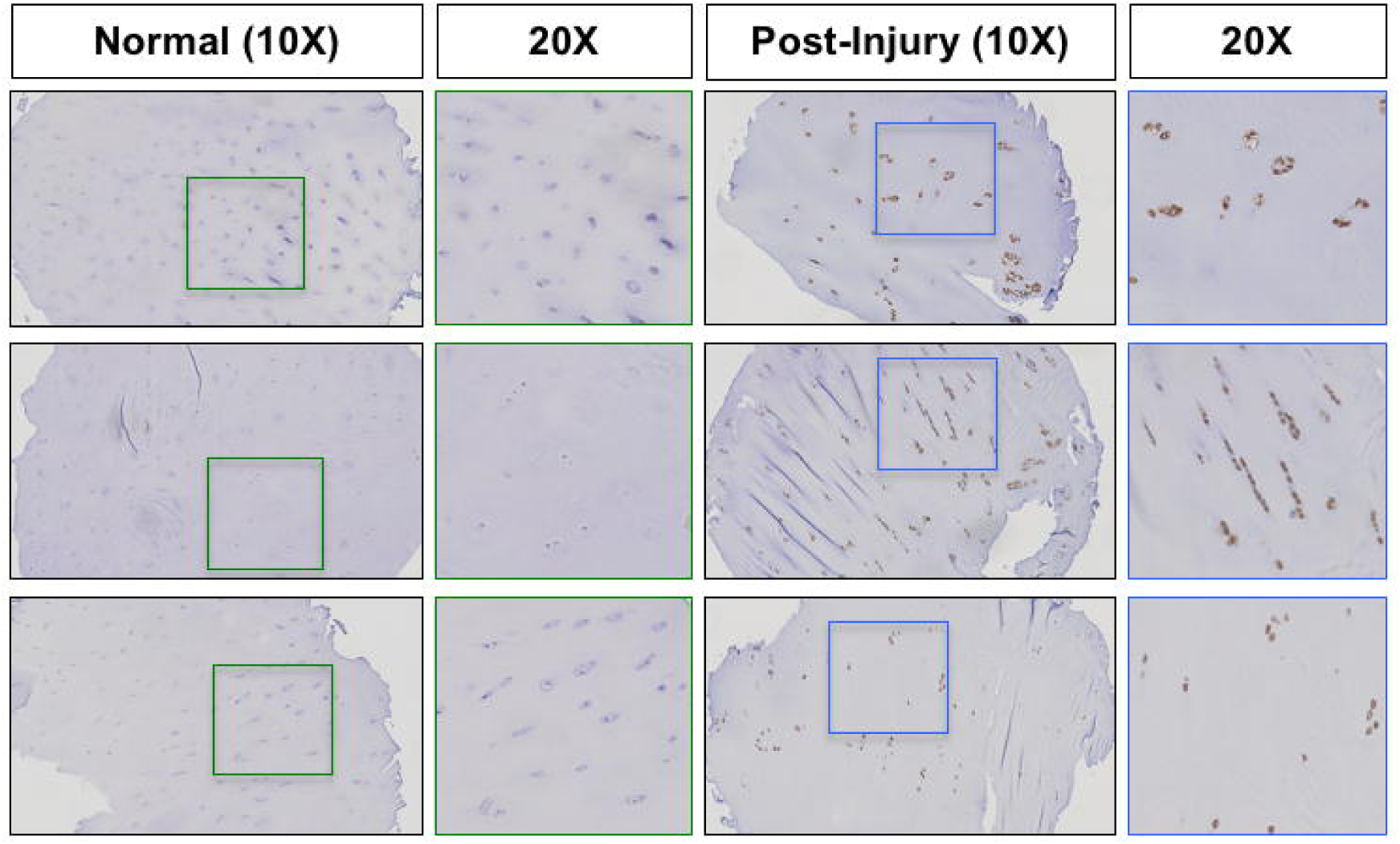
Knee joint injury leads to increased RUNX2 expression in human articular cartilage. Representative results of RUNX2 immunohistochemistry on a tissue microarray of normal human cartilage or cartilage from patients undergoing arthroscopic surgery 4 weeks following meniscal injury (Post-Injury). Left panels within each group are 10X images; right panels are 20X images of the corresponding boxed regions.

**Fig. 3.**
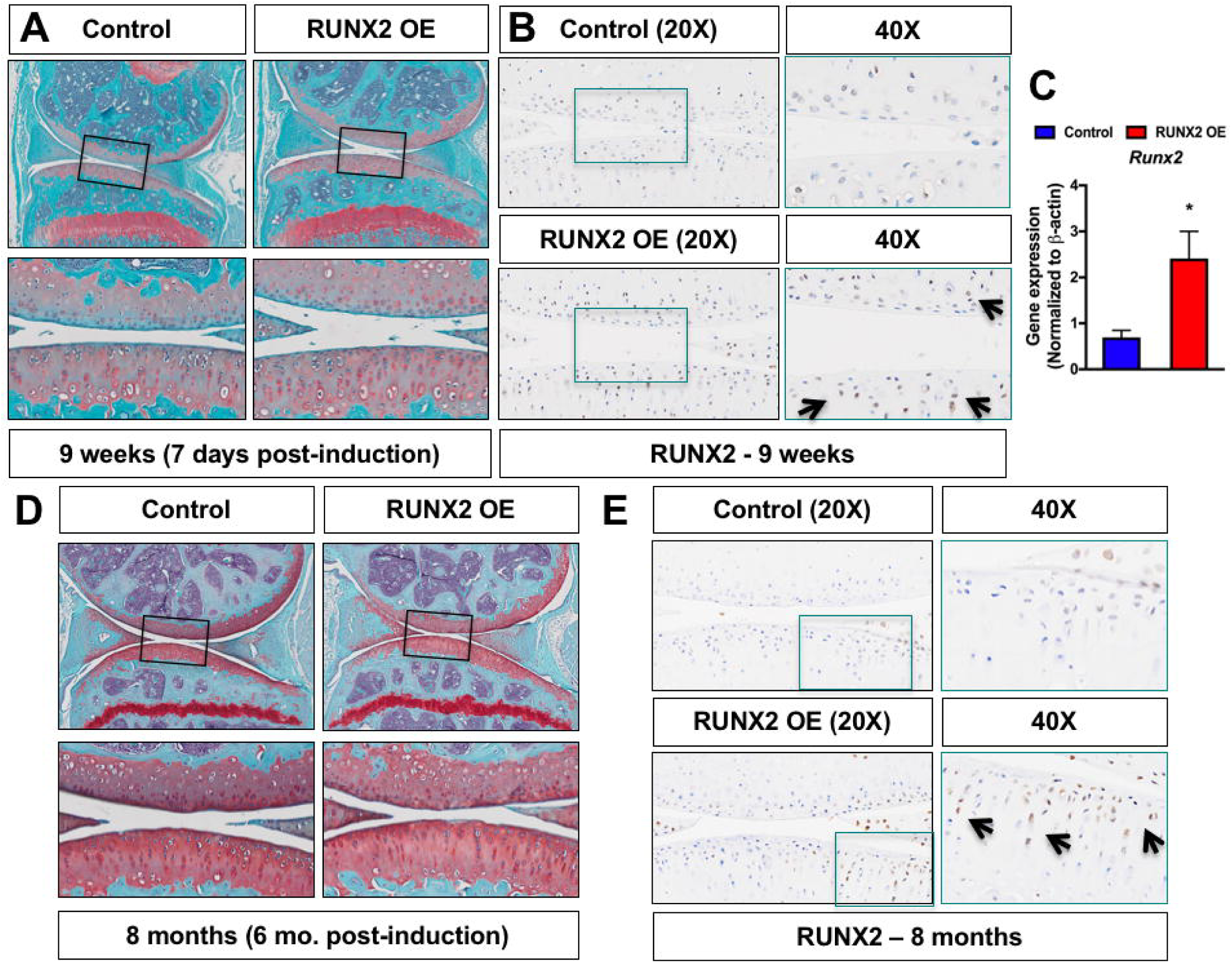
Chondrocyte-specific RUNX2 overexpression is insufficient to induce phenotypic changes in the articular cartilage. (A, D) Safranin O/Fast green staining of knee joint sections from male *R26^Runx2/+^* (Control) and *Acan^CreERT2/+^; R26^Runx2/+^* (RUNX2 OE) mice injected with tamoxifen at 2 months of age daily for 5 consecutive days and harvested 48 hours following the last tamoxifen injection at 9 weeks of age (A) or at 8 months of age (D). Top panels are 5X images of the knee joint; bottom panels are high magnification images (20X) of the boxed regions. (B, E) RUNX2 immunohistochemistry on knee joint sections from Control and RUNX2 OE mice at 9 weeks of age (B) and 8 months of age (E). Left panels are 20X images of the articular cartilage; right panels are high magnification images (40X) of the boxed regions. (C) Quantitative RT-PCR from tibial articular cartilage of Control and RUNX2 GOF mice harvested at 3 months of age following tamoxifen injections at 2 months of age (Control, n = 3, RUNX2 OE, n = 3). \**p* < 0.05, Student’s t-test.

### Postnatal chondrocyte-specific RUNX2 overexpression promotes accelerated articular cartilage degeneration following joint injury

It was shown previously that global haploinsufficiency or chondrocyte-specific loss of *Runx2* can inhibit the progression of post-traumatic OA following traumatic joint injury^(30,32)^. Since we did not find that RUNX2 OE alone in articular chondrocytes was sufficient to promote cartilage degeneration, however, we next tested whether chondrocyte-specific RUNX2 OE could affect the progression of OA in the injured joint environment. Specifically, we induced Cre-mediated recombination in both male and female Cre-negative control and RUNX2 OE (*Acan^CreERT2/+^; R26^Runx2/+^*) mice at 2 months of age using tamoxifen and subjected the mice to sham or MLI surgery at 2.5 months of age. Knee joints of control and RUNX2 OE mice were harvested 2 months following injury. Histologically, male RUNX2 OE mice showed enhanced articular cartilage degradation following MLI when compared to control mice, while sham sections did not reveal any detectable differences in the articular cartilage (**Fig. 4A**). To quantitatively assess the histological differences between male control and RUNX2 OE mice, modified OARSI scoring and manual histomorphometry were performed. MLI sections from RUNX2 OE mice showed significantly higher OARSI scores than those from control mice, and also showed reductions in overall tibial cartilage area (**Fig. 4B, C**). Tibial histomorphometric changes were seen in both the unmineralized and mineralized regions of the articular cartilage (**Fig. 4D**). SafO^+^ cartilage area was specifically reduced in the unmineralized tibial cartilage.

**Fig. 4.**
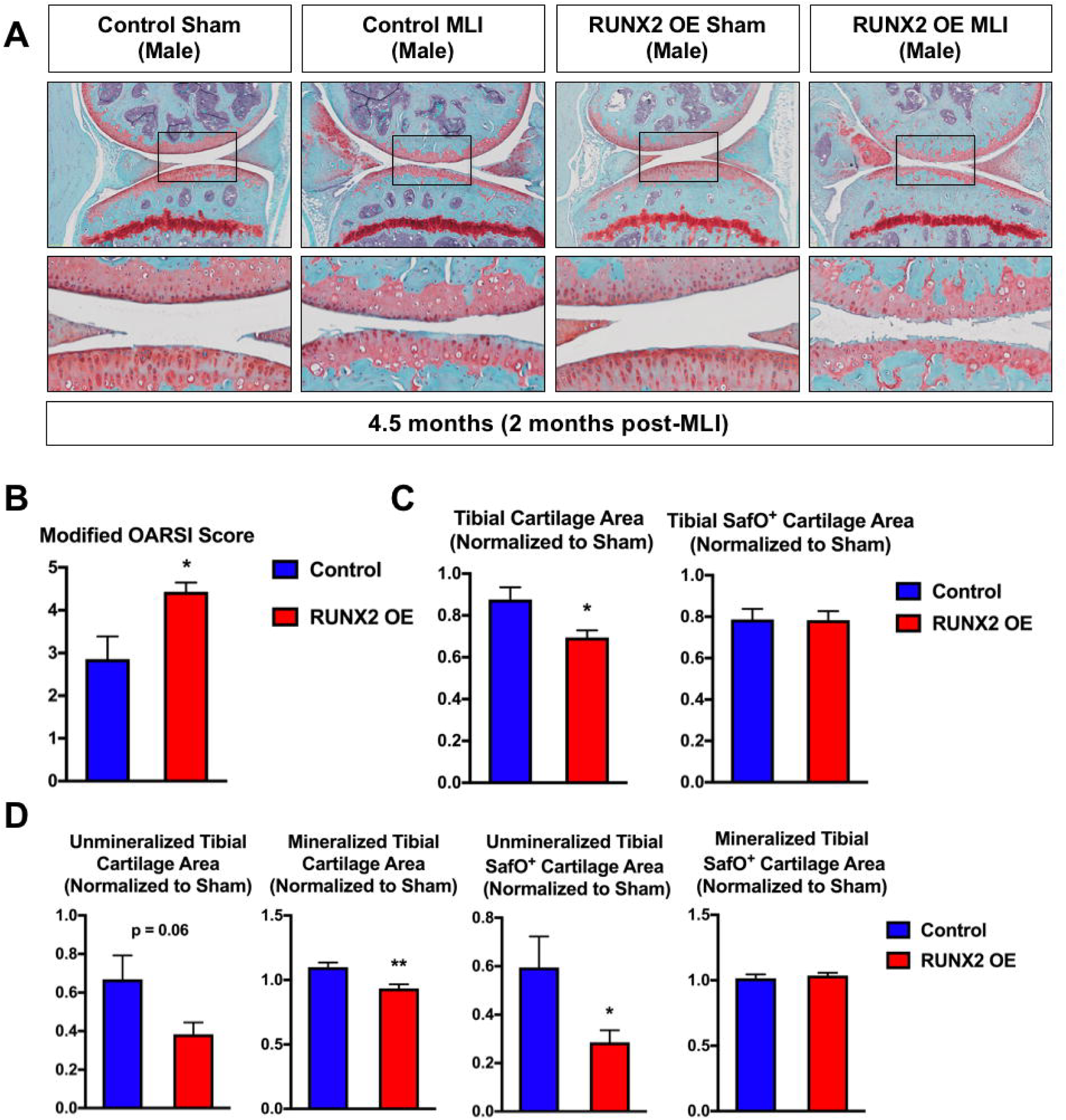
Postnatal RUNX2 overexpression accelerates articular cartilage degeneration following meniscal-ligamentous injury (MLI) in male mice. (A) Safranin O/Fast green staining of knee joint sections from male *R26^Runx2/+^* (Control) and *Acan^CreERT2/+^; R26^Runx2/+^* (RUNX2 OE) mice injected with tamoxifen at 2 months of age and subjected to sham or MLI at 2.5 months of age. Joints were harvested 2 months following injury. Top panels are 5X images of the knee joint; bottom panels are high magnification images (20X) of the boxed regions. (B) Modified OARSI scoring of Control and RUNX2 OE slides from the knee joint receiving MLI. (C, D) Quantitative histomorphometric analyses of total tibial cartilage area (C, left panel), total tibial SafO^+^ area (C, right panel), unmineralized and mineralized tibial cartilage areas (D, left panels), and unmineralized and mineralized tibial SafO^+^ areas (D, right panels) where MLI cartilage was normalized to corresponding sham cartilage in knee joint sections from Control (n = 6) and RUNX2 OE (n = 7) mice. \**p* < 0.05, \*\**p* < 0.01, Student’s t-test.

In contrast, female RUNX2 OE mice did not show differences in articular cartilage degradation in response to MLI relative to control mice (**Supplemental Fig. 3A**). This histological finding was confirmed by OARSI scoring and manual histomorphometry, which revealed no significant changes between the control and RUNX2 OE female mice following injury (**Supplemental Fig. 3B-D**).

In order to examine early molecular changes underlying the articular cartilage phenotype seen in the male RUNX2 OE mice 2 months following MLI, we performed a second set of sham or MLI surgeries on control and RUNX2 OE mice and harvested their knee joints one month following injury. Histology revealed decreased Safranin O staining within the tibial articular cartilage of the RUNX2 OE mice relative to controls, but these changes did not result in significant differences in modified OARSI score or manual histomorphometric measurements (**Fig. 5A, B**). We then performed immunohistochemistry for MMP13 and COL10A1, known downstream targets of RUNX2 in cartilage tissue^(24,26,46,47)^. Quantification of the MMP13 staining revealed a significant increase in MMP13^+^ chondrocytes present in the tibial articular cartilage of the RUNX2 OE mice (**Fig. 5D, E**). This increase in MMP13^+^ cells was primarily due to the increase in positive cells from the unmineralized region of the articular cartilage, as cell numbers were not significantly altered in the mineralized cartilage region (**Fig. 3D**). There was not a significant change in COL10A1-stained cartilage area in the RUNX2 OE mice.

**Fig. 5.**
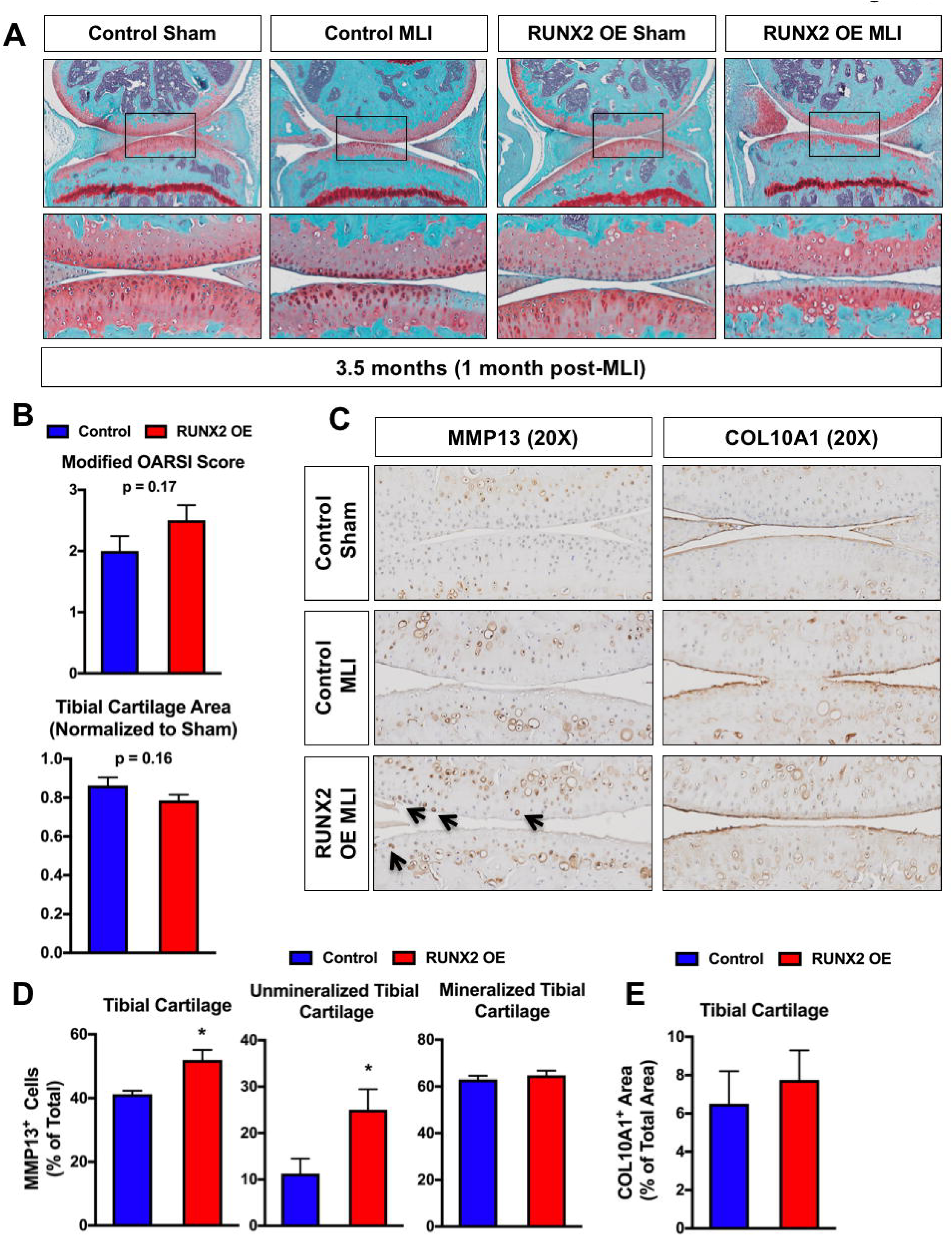
Postnatal RUNX2 overexpression induces MMP13 expression in unmineralized articular chondrocytes prior to accelerating articular cartilage degeneration following MLI. (A) Safranin O/Fast green staining of knee joint sections from *R26^Runx2/+^* (Control) and *Acan^CreERT2/+^; R26^Runx2/+^* (RUNX2 OE) mice injected with tamoxifen at 2 months of age and subjected to sham or MLI at 2.5 months of age. Joints were harvested one month following injury. Top panels are 5X images of the knee joint; bottom panels are high magnification images (20X) of the boxed regions. (B) Cartilage degeneration in Control (n = 6) and RUNX2 OE (n = 6) mice was evaluated by modified OARSI scoring (top graph) and quantitative histomorphometric analysis (bottom graph, data normalized to contralateral sham control sample). (C) MMP13 (left panels) and COL10A1 (right panels) immunohistochemistry of knee joint sections from Control or RUNX2 OE mice subjected to sham or MLI at 2.5 months of age and harvested 1 month following injury (20X images). (D) Quantification of MMP13^+^ cell number as percentage of total cell number from Control and RUNX2 OE mice (MLI samples only, n = 4 for both groups). (E) Quantification of COL10A1^+^ cartilage area as percentage of total cartilage area from Control and RUNX2 OE mice (MLI samples only, n = 4 for both groups). \**p* < 0.05, Student’s t-test.

Finally, given the increase in chondrocyte apoptosis due to RUNX2 OE during development, we next examined whether RUNX2 OE affected the amount of apoptotic cell death following traumatic joint injury. TUNEL staining was visibly increased within both the tibial and femoral articular cartilage of the RUNX2 OE mice relative to control mice following MLI and quantification of TUNEL^+^ cells relative to total articular chondrocyte numbers supported this observation (**Fig. 6A, B**).

**Fig. 6.**
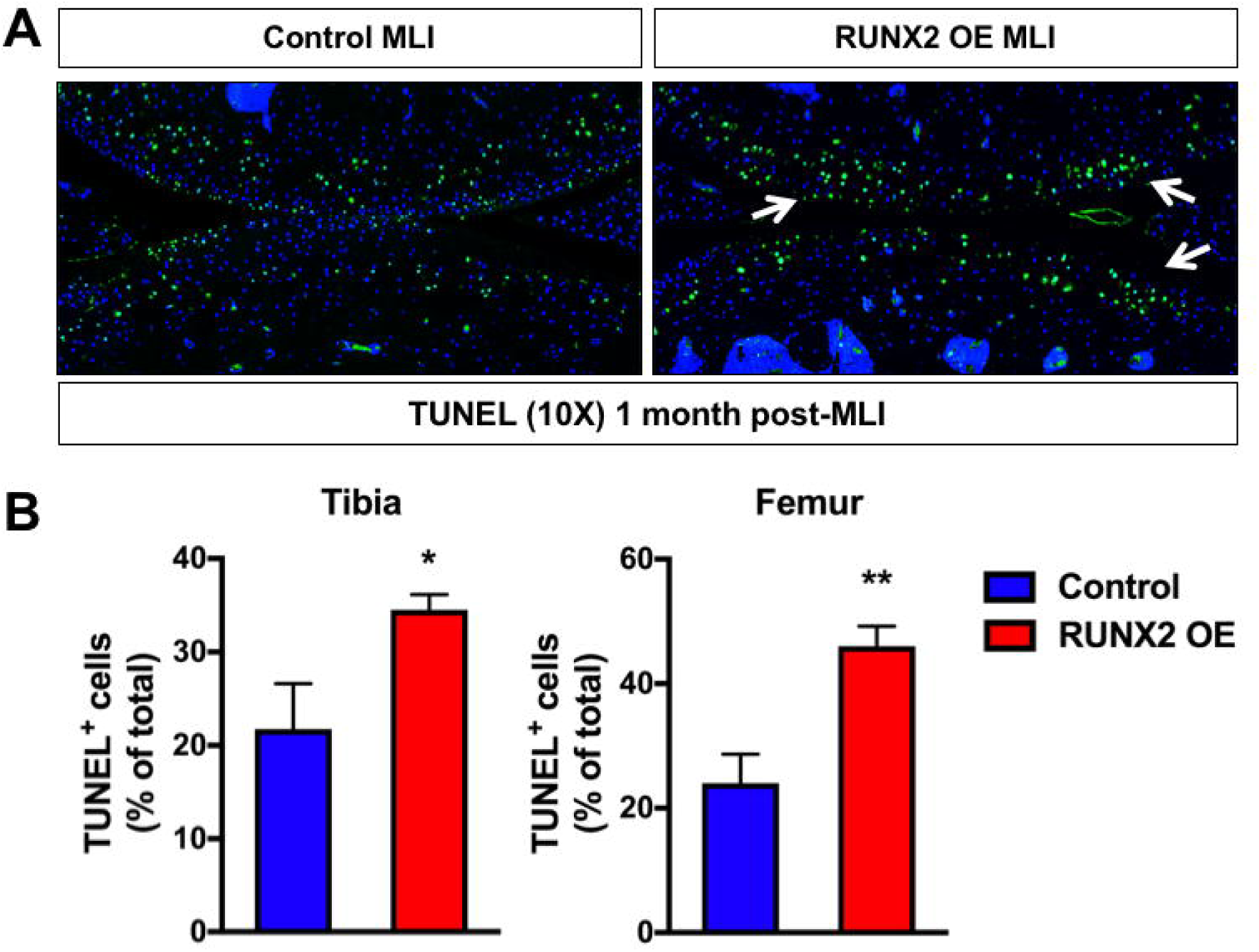
Postnatal RUNX2 overexpression results in enhanced articular chondrocyte apoptosis following meniscal-ligamentous injury (MLI). (A) TUNEL staining of knee joint sections from *R26^Runx2/+^* (Control) and *Acan^CreERT2/+^; R26^Runx2/+^* (RUNX2 OE) mice subjected to MLI at 2.5 months of age and harvested 1 month following injury (10X images). (B) Quantification of TUNEL^+^ cell number as percentage of total cell number from the tibial (left) or femoral (right) articular cartilage of Control or RUNX2 OE mice (MLI samples only, n = 4 for both groups). \**p* < 0.05, \*\**p* < 0.01, Student’s t-test.

## DISCUSSION

Our findings provide further confirmation that RUNX2 expression in growth plate chondrocytes during development promotes onset of hypertrophy and terminal maturation, resulting in chondrodysplasia. Previous models of exogenous RUNX2 expression in non-hypertrophic chondrocytes have relied on transgenic expression from the *Col2a1* promoter or expression of the *R26^Runx2^* allele in osteochondroprogenitor cells using a non-inducible *Col2a1-Cre*^(6,27,33)^. It is documented that *Col2a1* expression during development is not restricted to chondrocytes, but is first expressed in osteochondroprogenitor cells, those destined to become either chondrocytes or perichondrial pre-osteoblasts, prior to E12.5^(48-50)^. Therefore, it is possible that RUNX2 overexpression in this cell population may be at least partially responsible for the accelerated onset of hypertrophy observed in *Col2a1-Runx2* transgenic mice, as RUNX2 is known to drive FGF18 production in the perichondrium, inhibiting proliferation and promoting hypertrophy of neighboring chondrocytes^(51)^. Our model uses the *Col2a1-CreER^T2^* transgene to drive expression of the *R26^Runx2^* allele in a chondrocyte-specific manner by allowing induction of Cre-recombination at E13.5 ^(34,38,52)^. We found a slight, but significant, decrease in femur, tibia, and humerus length as well as disorganization of the proliferative and hypertrophic growth plate zones at E18.5 using this system. At E14.5, the hypertrophic zone and *Col10a1* expression domains were expanded and the *Mmp13* expression domain detectable in the *R26^Runx2^*-expressing embryos. At E18.5, the *Mmp13* expression domain appeared elongated at the chondro-osseous junction with increased expression in the trabecular bone, while the *Col10a1* expression domain was no longer expanded relative to total limb length. Collectively, these data suggest that RUNX2 expression in non-hypertrophic chondrocytes can accelerate onset of hypertrophy as well as late stages of chondrocyte maturation. The increased *Mmp13* expression in the trabecular bone is interesting and could be due to continued expression of exogenous RUNX2 in osteoblasts that are originally derived from targeted growth plate chondrocytes, as mounting evidence supports that hypertrophic chondrocytes can become osteoblasts during endochondral bone formation^(53-55)^.

Despite some differences in phenotype severity, the conclusions resulting from our developmental studies support those of the previous studies that investigated the effects of RUNX2 overexpression in non-hypertrophic chondrocytes. The differences in severity could be due to timing of induction, cell-type targeting specificity, or simply the level of exogenous RUNX2 produced. For example, Ueta, *et. al.* noted joint fusion and expanded regions of chondrocyte hypertrophy even at E18.5 in their *Col2a1-Runx2* transgenic mice, while we and others did not^(27)^. The *R26^Runx2^* allele was suggested by Tu *et. al.* to express a relatively low abundance of exogenous *Runx2* as they did not observe an increase in *Runx2* message by *in situ* hybridization with one or even two *R26^Runx2^* alleles^(33)^. They did observe, however, that the level of expression from the *R26^Runx2^* allele was sufficient to produce a functional protein product capable of rescuing bone formation in *Runx2^-/-^* embryos by promoting chondrocyte hypertrophy, perichondrial bone formation, and primary ossification center formation. In our developmental model, we were able to detect RUNX2 protein by IHC in some resting zone chondrocytes of the *C2T2; R26^Runx2^* and *C2T2; R26^Runx2/Runx2^* mutant mice. RUNX2 was not detected in this cell population in control mice consistent with the *Runx2* expression pattern reported by others^(4,6)^. We did not, however, see evidence of increased expression in pre-hypertrophic or hypertrophic chondrocytes suggesting that the level of expression above endogenous RUNX2 in these populations may be minimal. Nonetheless, the *R26^Runx2^* mouse provided the first opportunity to explore the role of RUNX2 in articular cartilage homeostasis and OA pathogenesis since other models resulted in early postnatal lethality or developmental defects precluding postnatal studies.

To our knowledge, this is the first study to assess the effects of exogenous RUNX2 expression in non-hypertrophic chondrocytes of adult mice and to test the hypothesis that expression of RUNX2 alone in this cell population is sufficient to promote the onset of OA. Despite detectable expression of RUNX2 mRNA and protein in the articular cartilage of mice immediately following induction and even 6 months later, we observed no difference in joint phenotype when compared to Cre-negative littermate control mice. Following surgical destabilization and meniscal injury of the joint, however, we found a significant difference in the rate of progression of post-traumatic OA in male mice. At 8 weeks following surgery, OARSI scores and histomorphometric measurements confirmed enhanced cartilage degeneration in RUNX2 OE mice. While there were no significant differences in OARSI scores or histomorphometric measurements 4 weeks following injury, we did find a significant increase in the number of MMP13^+^ and TUNEL^+^ cells suggesting increased cartilage catabolism and chondrocyte death in the injured RUNX2 OE joints. While *Mmp13* is a well-established direct downstream target gene of RUNX2^(23,24)^, it is less clear how RUNX2 might be affecting chondrocyte apoptosis. RUNX2 was shown to promote apoptosis in osteosarcoma cells via induction of *Bax* expression, but it has actually been shown to inhibit apoptosis in some other cell types^(44,56,57)^. Surprisingly, we did not find a significant difference in the amount of COL10A1 present in the cartilage matrix of the control and RUNX2 OE joints. We hypothesize that this might be due to decreased SOX9 expression in the injured joint^(58)^ (data not shown) and the requirement of SOX9 for *Col10a1* expression^(59)^.

Our studies included both male and female mice, but a significant difference in RUNX2-dependent cartilage degeneration following injury was only seen in male mice. Female mice, in general, develop less severe OA following traumatic knee joint injury than males and female-specific sex hormones offer protection against the development of post-traumatic OA, as ovariectomized female mice develop more severe OA following injury when compared to intact female mice^(60,61)^. Similarly, in humans, the loss of estrogen following the onset of menopause leads to a dramatic increase in the incidence of OA in women^(62,63)^. Estrogens have been shown to inhibit RUNX2 transcriptional activity in osteoblasts and breast cancer cells via the direct binding of estrogen receptor-α (ER-α) to the DNA-binding domain of RUNX2^(64)^. It may be possible, therefore, that RUNX2 activity is inhibited in the chondrocytes of the RUNX2 OE female mice despite elevated RUNX2 protein expression. ER-α is expressed in the articular chondrocytes of both mice and humans, but whether RUNX2 activity is altered by estrogen signaling in chondrocytes has yet to be determined^(65,66)^.

Due to the lack of a phenotype in the non-injured RUNX2 OE joints even in male mice, we hypothesize that the injured joint environment may potentiate RUNX2 activity and that RUNX2 activity is normally restricted in the articular chondrocytes of the unmineralized cartilage at homeostasis. RUNX2 is highly regulated through protein-protein interactions and post-translational modifications such as phosphorylation, acetylation, and ubiquitination to control its protein stability and/or transcriptional activity^(67-69)^. Signaling cascades activated by pro-inflammatory cytokines, oxidative stress, and mechanical stress all have the potential to affect RUNX2 activity within chondrocytes following joint injury. For example, p38 and ERK mitogen-activated protein kinases (MAPKs) are both known to positively regulate RUNX2 transcriptional activity in osteoblasts via direct phosphorylation and these kinases are activated in chondrocytes following cartilage injury^(70-75)^. While it is unknown whether they directly phosphorylate RUNX2 in chondrocytes, treatment of human articular chondrocytes with IL-1β leads to an induction of RUNX2-mediated MMP13 expression in a p38-dependent manner, suggesting that it may be possible^(76,77)^. GSK3β can also directly phosphorylate RUNX2 in osteoblasts, but on residues that inhibit transcriptional transactivation^(78)^. Further, treatment of chondrocyte cultures with GSK3β inhibitors was shown to promote nuclear localization of RUNX2 while inhibition of GSK3β *in vivo* enhanced cartilage degeneration in a murine model post-traumatic OA^(79,80)^. With regard to protein-protein interactions, RUNX2 binds to numerous other transcription factors, co-activators, and co-repressors that modulate its ability to affect gene expression^(69)^. C/EBPβ, for example, is a potent transcriptional partner of RUNX2 in regulation of *Mmp13* expression and is only expressed in articular chondrocytes following joint injury in a HIF-2α-dependent manner^(23)^. In contrast, the master chondrogenic transcription factor SOX9 was shown to promote RUNX2 degradation and to strongly suppress its activation through direct interaction^(81,82)^. While SOX9 is expressed in articular chondrocytes at homeostasis, as mentioned above, it is decreased in injured chondrocytes and in response to pro-inflammatory cytokine signaling^(58,83,84)^. Its loss following injury could, therefore, lead to activation of RUNX2 in the RUNX2 OE cartilage. Further studies, of course, will be required to determine if any of these or other potential mechanisms are responsible for the enhanced cartilage degeneration of the RUNX2 OE mice following joint injury.

Among the most important risk factors for the development OA in humans are prior traumatic joint injury and meniscectomy. It is estimated that approximately 12% of all symptomatic OA is related to prior injury and that 50% of all patients with an ACL or meniscus tear will develop knee OA^(44,45)^. A recent study comparing genetic risk variant loci in post-traumatic OA versus non-traumatic OA shows that the genetic contribution to the development of knee OA is just as high following injury as in primary, non-traumatic OA^(85)^. This suggests that underlying genetic variation may contribute to whether a patient will develop OA following injury and may even determine the rapidity of onset. Interestingly, large-scale genome-wide association studies (GWAS) have identified single nucleotide polymorphisms (SNPs) associating with OA near the *RUNX2* gene locus^(86)^. Additionally, hypomethylation of the *RUNX2* promoter was detected in a genome-wide methylation study comparing normal chondrocytes and OA chondrocytes^(87)^. Furthermore, OA candidate genes identified through GWAS studies include components of the WNT (*FRZB, DOT1L*), BMP (*GDF5*), and TGF-β (*SMAD3*) signaling pathways^(88-91)^. These pathways converge upon RUNX2 to regulate chondrocyte maturation. These findings suggest that genetic or epigenetic changes in *RUNX2*, or in the components of pathways that regulate RUNX2 expression and activity, may predispose to OA. Indeed, the genetic model we present here supports that RUNX2 overexpression in articular chondrocytes can accelerate the progression OA pathogenesis only following traumatic joint injury.

## ACKNOWLEDGEMENTS

This work was supported by grants from the National Institutes of Health/National Institute of Arthritis and Musculoskeletal and Skin Diseases (T32 AR053459, P30 AR061307, P30 AR069655, R21 AR07928 to JHJ and RE, and AR063071 to MJH). The authors thank Dr. Fanxin Long for his generous gift of the *R26^Runx2^* mouse. We also thank Center for Musculoskeletal Research Histology Core members Sarah Mack and Kathy Maltby for their technical assistance. Finally, we gratefully acknowledge the technical expertise of Dr. Erik Sampson in the creation of the human cartilage tissue microarray as well as Edward Ayoub and Tony Mirando for advice and technical assistance.

